# Role of trafficking protein particle complex 2 in medaka development

**DOI:** 10.1101/2023.07.14.548998

**Authors:** Francesca Zappa, Daniela Intartaglia, Andrea M. Guarino, Rossella De Cegli, Cathal Wilson, Francesco G. Salierno, Elena Polishchuk, Nicolina Cristina Sorrentino, Ivan Conte, Maria Antonietta De Matteis

**Affiliations:** Telethon Institute of Genetics and Medicine, TIGEM, Pozzuoli (Naples) Italy; Department of Molecular Medicine and Medical Biotechnology, University of Naples Federico II, Naples, Italy; Department of Biology, University of Naples Federico II, Naples, Italy; Department of Clinical Medicine and Surgery, University of Naples Federico II, Naples, Italy; present address: Altos Labs Bay Area Institute of Science, Altos Labs Inc., Redwood City (CA)

**Author notes:** These authors contributed equally to this work (Francesca Zappa, Daniela Intartaglia).

**Keywords:** medaka: Sedlin, spondyloepiphyseal dysplasia tarda, TRAPPC2, type II collagen

## Abstract

The skeletal dysplasia spondyloepiphyseal dysplasia tarda (SEDT) is caused by mutations in the *TRAPPC2* gene, which encodes Sedlin, a component of the trafficking protein particle (TRAPP) complex that we have shown previously to be required for the export of type II collagen (Col2) from the endoplasmic reticulum. No vertebrate model for SEDT has been generated thus far. To address this gap, we generated a Sedlin knockout animal by mutating the orthologous *TRAPPC2* gene (*olSedl*) of *Oryzias latipes* (medaka) fish. *OlSedl* deficiency leads to embryonic defects, short size, diminished skeletal ossification, and altered Col2 production and secretion, resembling human defects observed in SEDT patients. Moreover, SEDT knock-out animals display photoreceptor degeneration and gut morphogenesis defects, suggesting a key role for Sedlin in the development of these organs. Thus, by studying Sedlin function *in vivo*, we provide evidence for a mechanistic link between TRAPPC2-mediated membrane trafficking, Col2 export, and developmental disorders.

## Introduction

Type II collagen, encoded by the *COL2A1* gene, is the major component of cartilage. *COL2A1* mutations cause type II collagenopathies (OMIM#120140) which include diverse clinical entities such as achondrogenesis (OMIM#200610), Stickler syndrome (OMIM#108300), and spondyloepiphyseal dysplasia congenita, (SEDC, OMIM#183900). Common manifestations of such diseases are impaired chondrogenesis and osteogenesis that vary in severity depending on the mutation. Mutations of *COL2A1*, which is abundantly expressed in the vitreous (Jobling *et al,* 2014), can also lead to extraskeletal manifestations, the most common involving defects ranging from myopia to severe vitreoretinal degeneration (Jabobson *et al,* 2023).

A distinct form of skeletal dysplasia called spondyloepiphyseal dysplasia tarda (SEDT, OMIM#313400) is caused by mutations in the *TRAPPC2* gene. SEDT is less severe than SEDC, arises in late childhood, and is characterized by short stature, disproportionate short trunk, and precocious mild osteoarthritis, which manifests in the early teens (Whyte *et al,* 1999). The *TRAPPC2* (or *SEDL*) gene is located on chromosome Xp22.2 and encodes the ubiquitously expressed Sedlin/TRAPPC2 protein (Gedeon *et al*, 1999). Sedlin is a core component of the highly conserved oligomeric TRAnsport Protein Particle (TRAPP) complex that controls different segments of vesicular transport (Sacher *et al,* 2019). The mammalian TRAPP complex exists in two different forms, TRAPPII and TRAPPIII. Both complexes, which control different stages of trafficking, have common core components but differ in their peripheral elements. Sedlin functions as a connector between the shared core and specific peripheral subunits.

We have shown that Sedlin, in addition to interacting with other TRAPP components, interacts with the small GTPase Sar1, modulating its GTP-GDP cycle by promoting GTP hydrolysis, a key event for the formation of ER-derived carriers (Venditti *et al*, 2012). Due to this activity and the ability to interact with the procollagen export factor TANGO1 at ER-exit sites (ERES), Sedlin is required for efficient procollagen export, as shown in cultured chondrocytes and in SEDT patient fibroblasts (Venditti *et al*, 2012).

TRAPPC2 is evolutionarily conserved, ubiquitously expressed in mammals, and essential in yeast (Gedeon *et al*, 1999; Gécz *et al*, 2003). The cartilage and bone restricted phenotype in SEDT patients seem contradictory but could be explained by the existence of an expressed *TRAPPC2* pseudogene on chromosome 19. This pseudogene encodes a protein identical to Sedlin, potentially compensating for the absence of the protein encoded by the gene on the X chromosome (Gécz *et al*, 2000; Ghosh *et al*, 2001; Vinckenbosch *et al*, 2006). Notably, the TRAPPC2 pseudogene is also present in mice (0610009B22Rik) and in rats (Gene ID: 100910318), and to date no functional studies have been performed *in vivo* to elucidate the role of Sedlin. We therefore decided to address the *in vivo* role of TRAPPC2 in a well-characterized teleost model for the study of osteochondral diseases, Japanese medaka (*Oryzias latipes*) (Lin *et al,* 2016), which has only one copy of the *TRAPPC2*/*SEDL* gene, hereon called *olSedl*.

We show that *olSedl* deletion induces a skeletal phenotype reminiscent of SEDT, but also extraskeletal defects in the eye and gut. The marked defects in collagen deposition that we find in the Sedlin-deficient fish may provide the common basis for skeletal and extraskeletal manifestations, as suggested by the co-occurrence of eye involvement in syndromic forms of osteodysplasia caused by COL2 mutations (Jobling *et al,* 2014) and by the known role of the extracellular matrix (ECM), including collagen, in intestinal epithelium polarization (Pompili *et al,* 2021). Our data suggest that Sedlin is required for vertebrate embryo development by regulating collagen production and secretion in different tissues.

## Results

### Generation of Sedlin knock-out medaka fish

The medaka genome harbours a single *olSedl* gene (http://www.ensembl.org/Oryzias_latipes; ENSORLG00000025160) that encodes a protein with 90% homology to human Sedlin (Fig EV1A). Quantitative real-time PCR (qRT-PCR) analysis revealed high levels of *olSedl* expression from stage (st) 32 to 37 (Fig 1A) when organogenesis and bone tissue mineralization are occurring in medaka embryos (Iwamatsu, 2004; Kupsco and Schlenk 2016). Notably, *olSedl* expression paralleled that of *TRAPPC3*, another core component of the TRAPP complex, and of *Col2a1a* (Fig 1A). Whole-mount in situ hybridization (ISH) using a DIG-labelled antisense RNA probe showed ubiquitous *olSedl* expression, including in vertebrae and trunk muscle of larvae at stage 40 (Fig EV1B).

**Figure 1.**
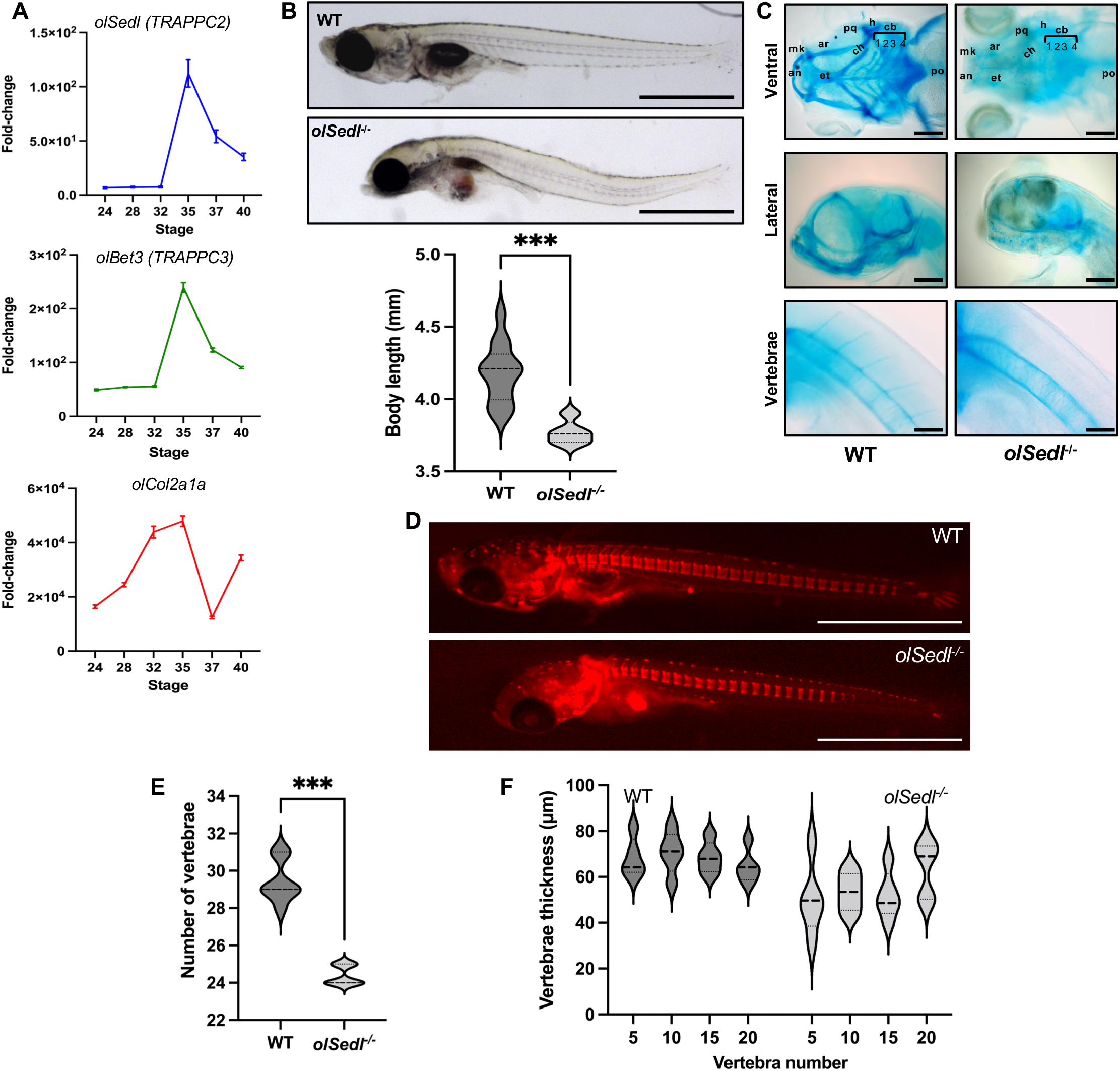
*olSedl*^-/-^ mutants display a SEDT-like phenotype. A. RT-qPCR analysis of *olSedl* (*TRAPPC2*), *olBet3* (*TRAPPC3*) and *olCol2a1a* expression at the indicated developmental stages. Values are expressed as fold change relative to stage 19. B. Representative brightfield images of stage 40 WT and *olSedl*^-/-^ individuals showing that *olSedlin* depletion induces evident trunk curvature and reduced body length. Violin plots showing body length quantification are reported below. C. Representative images of craniofacial alcian blue staining (from top to bottom: ventral plane, lateral plane, and vertebra detail) showing a marked reduction of cartilaginous intermediates in *olSedl*^-/-^ fish [(an) anterior limit, (ar) articulation, (cb1-4) ceratobranchial pairs 1 to 4, (ch) ceratohyal, (et) ethmoid plate, (h) hyosymplectic, (mk) Meckel’s cartilage, (pq) palatoquadrate, (po) posterior limit]. D. Representative Alizarin Red S vital staining images showing a reduction in the number of calcified vertebrae in stage 40 *olSedl*^-/-^ individuals. E. Violin plots showing quantification of the number of vertebrae per fish. F. Vertebrae thickness measurements for WT and *olSedl*^-/-^ fish. Data information: the dashed line and punctate pattern in the violin plots in (B) (E) and (F) show the median and quartiles, respectively. ***p < 0.001, two-tailed unpaired t-test with Welch’s correction (B), **p = 0.002, two-tailed unpaired Mann-Whitney test (E). (B, E, F): WT N = 14, *olSedl^-/-^* N = 5. Scale bars in (B) and (D): 1 mm; (C): 250 μm.

Prompted by these results, we used Transcription activator-like effector nuclease (TALEN)-targeted mutagenesis (Ansai *et al,* 2013) to generate an *olSedl* knockout (KO) medaka line (Fig EV1). Four different mutations were isolated including sense and nonsense deletions with a total targeting efficiency of 16.7%. Three of the mutations *(olSedl*^Δ101-109/Δ101-109^, *olSedl*^Δ113-117/Δ113-117^ and *olSedl*^Δ111-118/Δ111-118^) induced a frameshift and a premature termination of the protein, reminiscent of mutations found in SEDT patients (Zhang *et al,* 2020). Of these three *olSedl* KOs, we selected *olSedl*^Δ111-118/Δ111-118^ (hereon *olSedl^-/-^*) (Figs EV1C-E) for further analysis.

### *OlSedl* is required for skeletogenesis

No severe morphological abnormalities or defects in somatogenesis were observed in *olSedl^-/-^* compared to wild-type embryos during the initial stages of development. However, *olSedl^-/-^* larvae at stage 40 displayed a delay in mean hatching time, suggesting a severe reduction of prenatal motility. Indeed, *olSedl^-/-^* showed a dramatic reduction in larval movement and died within the first few hours after hatching (Movies EV1 and EV2).

*olSedl^-/-^* larvae also exhibited abnormalities in craniofacial morphogenesis, a shorter body, and a slight curvature of the trunk in the dorsal-ventral and medial-lateral planes, accompanied by highly penetrant bone defects (Fig 1B). Cartilaginous elements in the head, including the ceratobranchial pairs, ceratohyal, and ethmoid plate, were almost absent in *olSedl^-/-^* fish, leading to a completely dysmorphic craniofacial skeleton compared to control fish (Fig 1C). As a consequence, *olSedl^-/-^* displayed an evident decrease in bone mineralization of the head, caudal fin rays, and skeletal vertebrae (Fig 1D).

When analysed by alizarin red staining, the number of calcified vertebrae at st40 was significantly lower than in WT animals (-18.5% ± 1.2 SD, Fig 1E). Furthermore, a reduction in the vertebrae thickness was detected in *olSedl^-/-^* (Fig 1F), in line with the spine phenotype observed in SEDT patients (Ruyani *et al,* 2012).

To assess whether, as observed in cell systems, Sedlin controls collagen deposition also *in vivo*, we performed whole-mount immunofluorescence analysis for Col2A on larvae at st40. A notable reduction of Col2A in the ECM was found in *olSedl^-/-^* larvae (Fig 2A), which correlates with a reduction of protein levels (Fig EV1E) and with diminished ECM thickness (Fig 2B).

**Figure 2.**
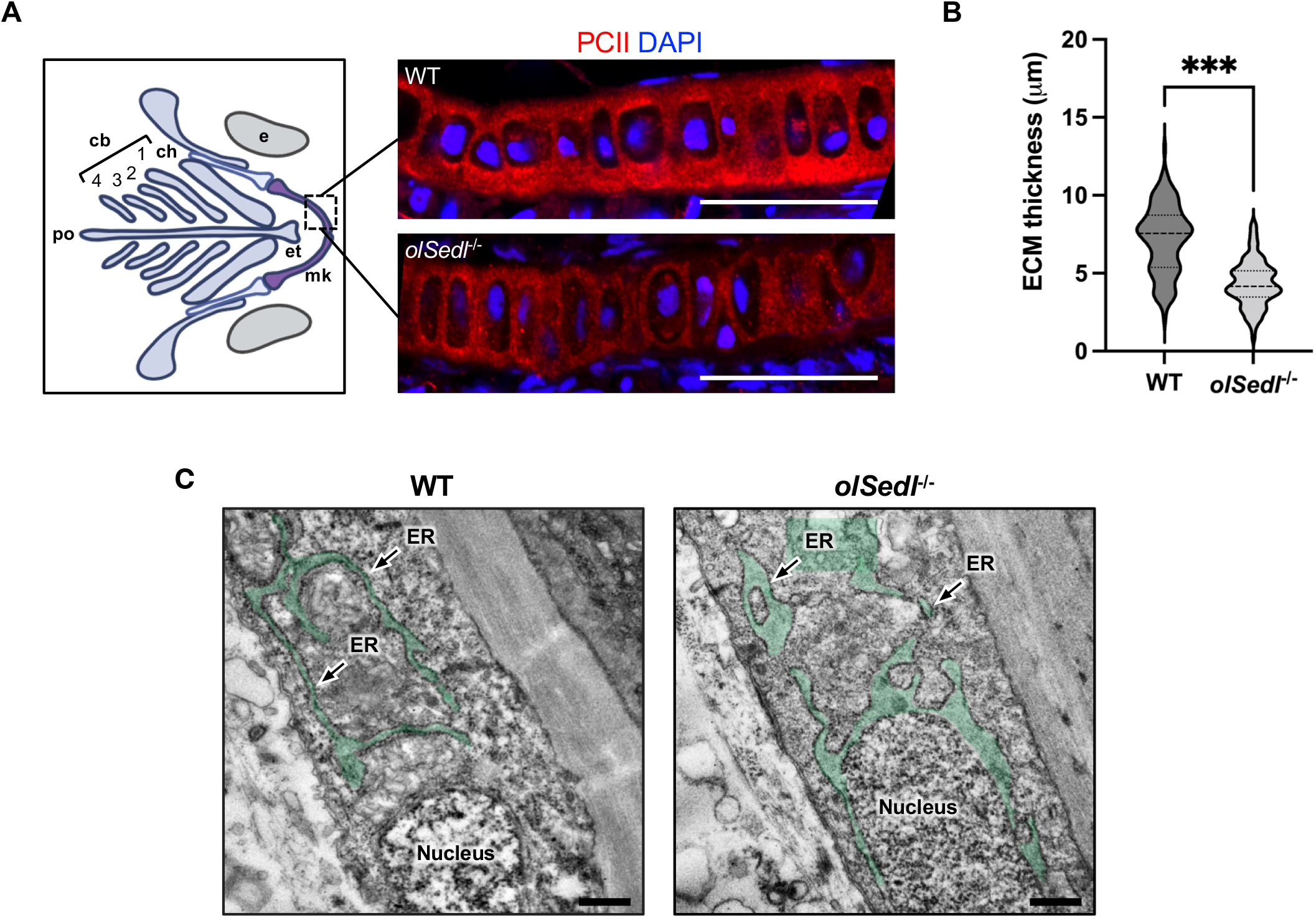
Col2 expression and secretion is compromised in *olSedl*^-/-^ fish. A. Representative immunostaining of Meckel’s cartilage (boxed area in the graphic representation on the left) of WT and *olSedl*^-/-^ fish showing a reduction in type II collagen (red) levels. Nuclei are counterstained with DAPI (blue) [(cb1-4) ceratobranchial pairs 1 to 4, (ch) ceratohyal, (et) ethmoid plate, (mk) Meckel’s cartilage, (po) posterior limit, (e) eye]. B. Quantification of ECM thickness in WT and *olSedl*^-/-^ fish. C. Representative electron microscopy images (16x magnification) of a vertebral section of stage 40 WT and *olSedl*^-/-^ fish. The ER is pseudocoloured in green and the ER cisternae are indicated by black arrows. Data information: the dashed line and punctate pattern in the violin plot in (B) show the median and quartiles, respectively. (B): n > 90 measurements. ***p < 0.001, two-tailed unpaired t-test with Welch’s correction. Scale bar in (A): 30 μm, (C): 500 nm.

Transmission Electron Microscopy (TEM) analysis performed on larvae vertebrae revealed ER expansion in *olSedl^-/-^* cells (Fig 2C). Of note, fibroblasts derived from SEDT patients show ER dilation and acute Sedlin depletion in cell models leads to ER accumulation of Col2A and consequent ER expansion (Venditti *et al,* 2012).

To rule out possible TALEN off-target effects (Koo *et al,* 2015), we injected WT fertilized eggs with a specific morpholino (MO) directed against the Sedlin ATG initiation codon within the 5′ untranslated region (MO_Sedl_) (Robu *et al,* 2007; Eisen & Smith, 2008) (Fig 3A). MO_Sedl_ display skeletal defects (70 ± 5% of 3,000 injected embryos) including trunk curvature, reduction of cartilaginous elements, i.e. vertebrae and caudal fin rays, and absence or reduction of the neural arch (Fig 3B-E). Moreover, MO_Sedl_ show a reduction of Col2A, phenocopying *olSedl^-/-^* and thus corroborating our findings (Fig 3F).

**Figure 3.**
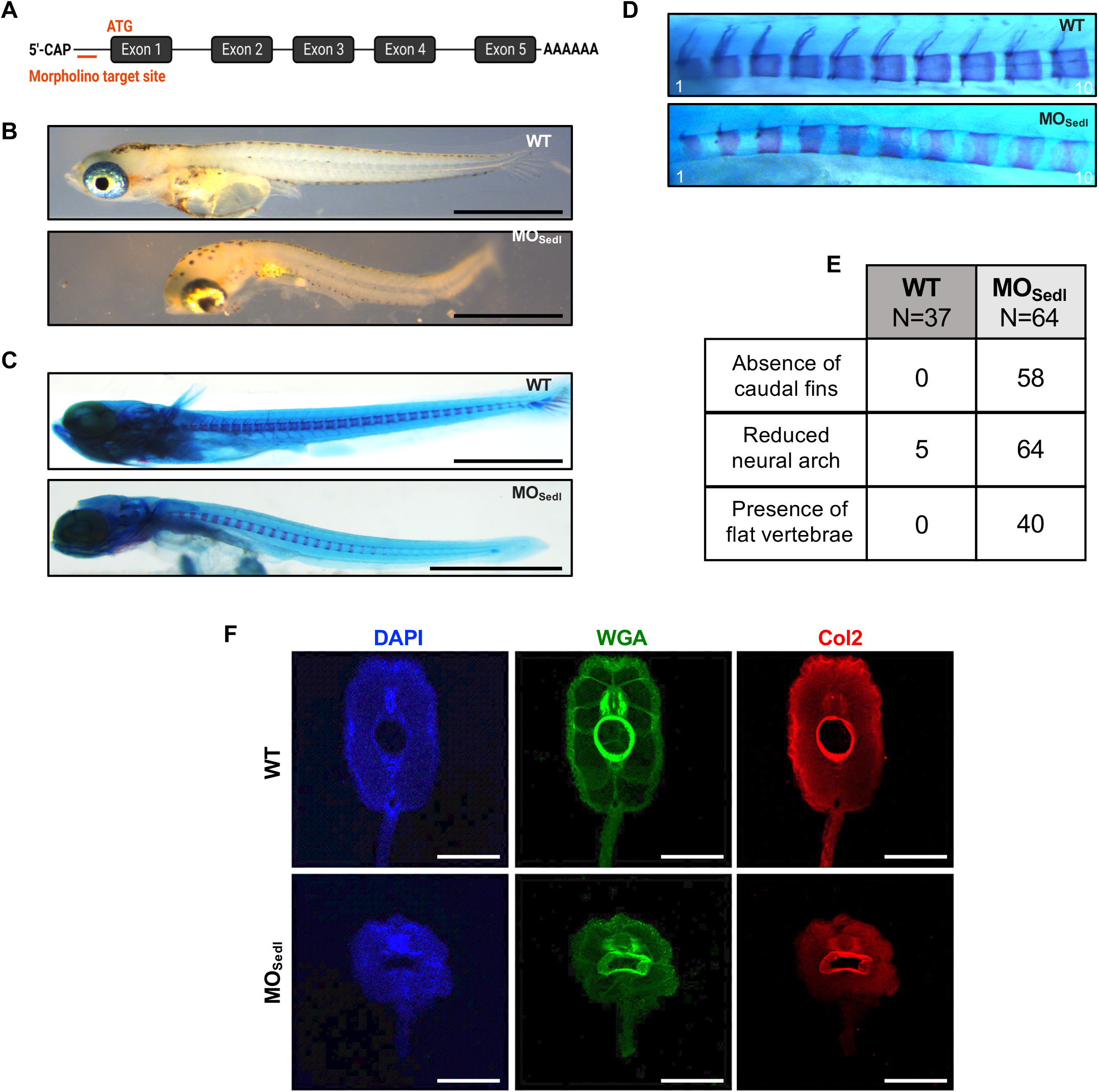
*olSedl* morpholino fish manifest a SEDT-like phenotype. A. Schematic representation of the morpholino targeting site at the ATG in exon 1 of *olSedl*. B. Representative brightfield images of WT and MO_Sedl_ fish. C. Representative whole-mount alcian blue-alizarin red staining images showing that MO_Sedl_ lack cartilaginous intermediates in the caudal fin and have a reduced number of vertebrae. D. High magnification alcian blue-alizarin red staining from the 1^st^ to 10^th^ vertebra showing altered morphology. E. Quantification of the observed phenotypes. F. Immunofluorescence analysis of WT and MO_Sedl_ sections of frontal vertebra showing a significant alteration in size and morphology. Nuclei are counterstained with DAPI (blue), connective tissue is visualised with fluorescent Wheat Germ Agglutinin (WGA, green), while type II collagen (Col2) is shown in red. Data information: Scale bar in (B) and (C): 1 mm; (F): 100 μm.

These data indicate that *olSedl^-/-^* recapitulates the major clinical signs observed in humans with TRAPPC2 mutations and represents a suitable model to study SEDT.

### Transcriptome changes induced by Sedlin depletion

To gain an unbiased view of the impact of Sedlin depletion on medaka fish development, we performed transcriptomics analysis on both morpholino-induced MO_Sedl_ and on *olSedl^-/-^*embryos to mitigate the risk of transcriptional changes not directly or specifically due to the absence of functional Sedlin.

QuantSeq 3’mRNA-Seq analysis was performed on *olSedl^-/-^*embryos while RNA-Seq was performed on MO_Sedl_ embryos (Fig EV2). QuantSeq 3’mRNA-Seq (GEO accession GSE143538) revealed 1,052 differentially expressed genes (414 genes induced and 638 inhibited), while 4,005 differentially expressed genes (1,870 genes induced and 2,135 inhibited) were found by RNA-Seq (GEO accession GSE186769) (Fig EV2A). Differentially Expressed Genes (DEGs) were converted into the corresponding human orthologues using the BioMart browser (Cunningham *et al,* 2022).

We found 695 DEGs in common between the two datasets, of which 254 were induced and 399 were inhibited genes (Fig EV2A). This comparison of the MO_Sedl_ and *olSedl^-/-^* data increased the confidence that the changes were due specifically to the loss of Sedlin.

Among the most down-regulated gene classes were genes involved in extracellular matrix organization (GO:0030198), but also genes involved in visual perception (GO:0007601) and intestinal absorption (GO:0050892) (Figs EV2B, EV3). On the one hand, these findings are consistent with the observed skeletal phenotype (by highlighting a dysregulation of ECM components) while on the other they point to a possible role of Sedlin in controlling eye and intestine development.

### *olSedl* oversees chondrocyte differentiation

Collagens and collagen-related genes were among the most affected of the down-regulated genes (Fig 4A and Fig EV2C) with the *col2a1* gene being one of the most inhibited genes in the *olSedl^-/-^*. The expression of type XI collagens (*col11a1* and *col11a2*), which are part of the nucleating core of type II collagen fibrils, and of *col10a,* a marker for chondrocyte maturation (Maye *et al,* 2011), was also significantly reduced in *olSedl^-/-^*(Fig 4A). The impaired Col10 expression was validated in medaka lines expressing GFP under the control of the *col10a1* promoter (Renn *et al,* 2013). Here, GFP fluorescence was significantly reduced in *olSedl^+/-^* heterozygous fish and almost lost in *olSedl^-/-^*(Fig 4B).

**Figure 4.**
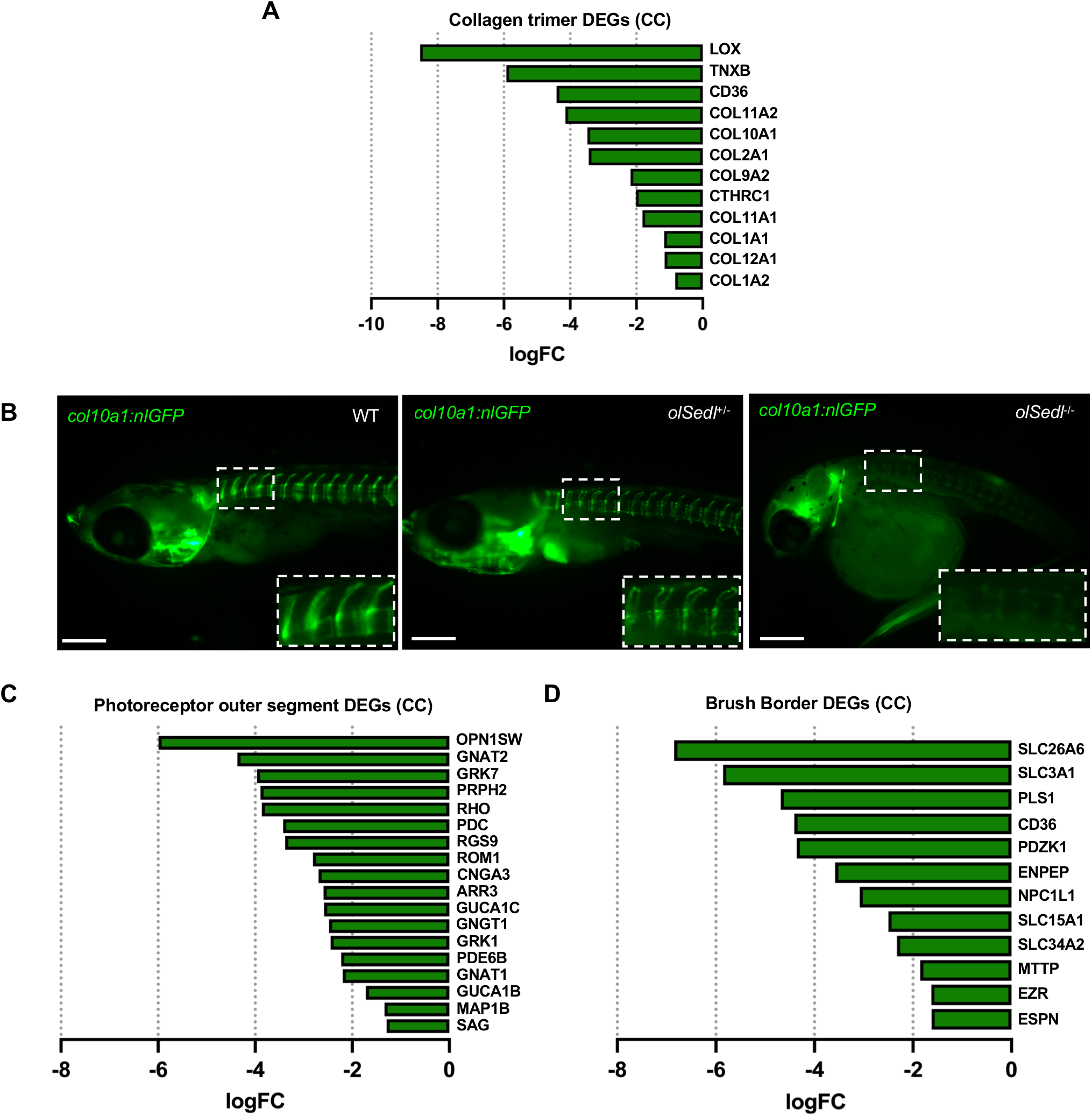
Sedlin depletion affects expression of genes involved in skeletogenesis, retinogenesis, and intestinal epithelium development. A. Graphical representation of genes in the CC (Cellular Compartments) class *GO:0005581∼collagen trimer* significantly downregulated in the QuantSeq-3’-mRNA-Seq dataset. B. Representative images of WT, heterozygous (*olSedl*^+/-^), and homozygous (*olSedl*^-/-^) fish, transgenic for *col10a1:nlGFP.* Insets, enlarged images of boxed areas showing GFP signal in vertebrae. Scale bar, 100 μm. C. Graphical representation of genes in the CC class *GO:0001750∼photoreceptor outer segment* significantly downregulated in the QuantSeq-3’-mRNA-Seq dataset. E. Graphical representation of genes in the CC class *GO:0005903∼brush border* significantly downregulated in the QuantSeq-3’-mRNA-Seq dataset. See Fig EV3 for downregulated genes in both QuantSeq-3’-mRNA-Seq and RNA-Seq datasets.

One possibility for the reduction in Col2A mRNA expression was that transcription factors controlling its expression might be affected by Sedlin depletion. However, we found no significant changes in the expression levels of the main transcription factors upstream of *col2a1,* such as *Sox9b*, *Sox5,* and *Sox6* (Kozhemyakina *et al,* 2015), in *olSedl^-/-^*. It has been reported that altered ECM due to Col6A2-KO in mice leads to abnormal chondrocyte differentiation and maturation associated with altered ECM-gene expression profiles (Komori *et al,* 2022) and that the absence of aggrecan affects the co-expression of genes encoding collagens II, X, and XI (Wai *et al,* 1998). Thus, alteration of one ECM component may lead to alterations in the expression of other ECM components that together contribute to the observed phenotype, but how this occurs mechanistically is still unclear. This appears to be the case here, where impairment of Col2A secretion due to the lack of Sedlin causes ECM remodeling and induces a feed-forward loop that negatively impacts on the expression of ECM genes.

### Sedlin is required for photoreceptor development

As mentioned above, our transcriptomics dataset revealed that Sedlin depletion strongly downregulated genes involved in visual perception (GO:0007601), detection of light stimulus (GO:0009583), and visible light (GO:0009584). For example, the expression of photoreceptor markers such as GNAT1, GNAT2 and GNGT1, encoding the Transducin 1,2 and gamma subunits which couple rhodopsin stimulation and cGMP-phosphodiesterase (Lagman *et al,* 2015) (Fig 4C), and RPGRIP1 (logFC, -3.2), encoding a photoreceptor component of cone and rod photoreceptor cells (Mavlyutov *et al,* 2002), was strongly impaired.

Similarly, the expression of genes involved in the maintenance of retinal integrity such as ROM1, a member of a photoreceptor-specific gene family that encodes Rod Outer Segment Protein, an integral membrane protein found in the photoreceptor disk rim of the eye (Conley *et al,* 2019) (Fig 4C), and RS1 (logFC, -2.915), encoding retinoschisin, an extracellular protein that plays a crucial role in the cellular organization of the retina (Molday *et al,* 2007), was also reduced.

The reduced expression of genes involved in signal transduction and retinal integrity prompted us to explore the biological relevance of these results, evaluating whether *olSedl* ablation affected photoreceptor differentiation and maintenance during eye development.

We found that while the lamination of the retina was preserved, retinas from *olSedl^-/-^*displayed significant alterations in photoreceptor marker staining, which was more evident in ventral regions compared to WT larvae (Fig 5). Specifically, a strong reduction of rhodopsin-positive mature rod photoreceptors was accompanied by a decrease in Zpr-1-positive mature cone photoreceptors (Fig 5A and 5B). Notably, outer/inner segments of both cones and rods appeared significantly shorter and morphologically abnormal in *olSedl^-/-^* retina compared to WT siblings (Fig 5A and 5B). The disorganization of the photoreceptors detected at st40 was already visible at st36 (Fig EV4A and EV4B), supporting an alteration of photoreceptor development. Consistent with defective photoreceptor differentiation and structures, TEM analysis confirmed that photoreceptor outer segments were shorter and thinner in *olSedl^-/-^* retinas (Fig 5C and 5D). However, we did not detect any apparent morphological defects in the retinal pigment epithelium of *olSedl^-/-^* with respect to WT controls, as determined by TEM analysis.

**Figure 5.**
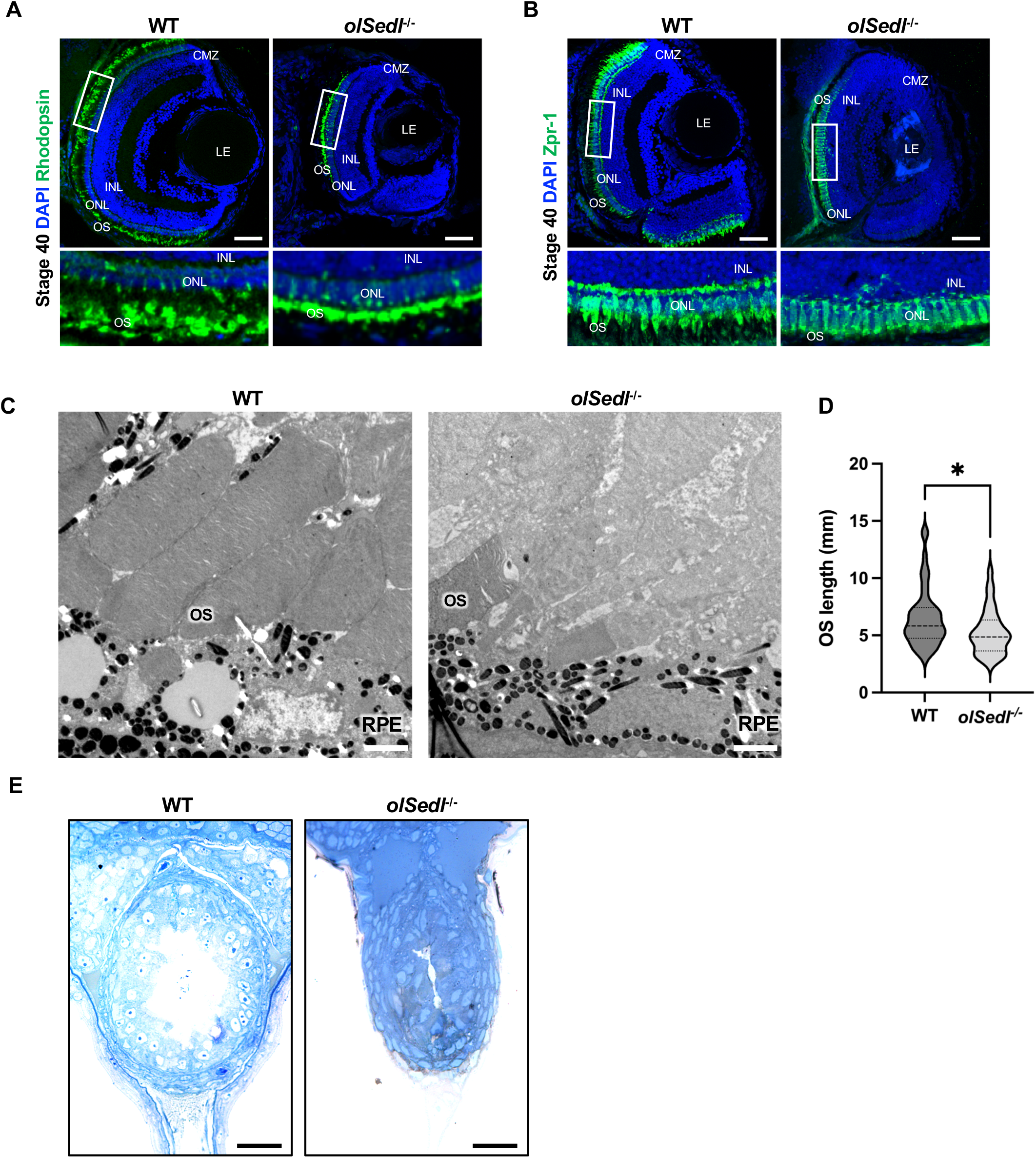
*olSedl^-/-^* fish manifest altered photoreceptors and gut epithelium. A, B. Representative images of Rhodopsin (A, green) and Zpr-1 (B, green) immunostaining of frozen retinal sections of WT and *olSedl^-/-^* stage 40 individuals. Nuclei are counterstained with DAPI (blue). Lower panels are enlarged images of the boxed areas showing discrete Rhodopsin and Zpr-1 staining in the rod and cones of the Outer Segment (OS) layer. *olSedl^-/-^* fish show altered ONL/INL patterning [(CMZ) ciliary marginal zone, (OS) outer segment, (ONL) outer nuclear layer, (INL) inner nuclear layer, (LE) lens]. C. Representative Transmission Electron Microscopy (TEM) images of the OS of WT and *olSedl^-/-^* fish. Photoreceptor cells of *olSedl^-/-^*fish are shorter and thinner than WT [(OS) outer segment, (RPE) retinal pigment epithelium]. D. Quantification of the length of the outer segment. The dashed line and punctate pattern in the violin plot show the median and quartiles, respectively. E. Toluidine blue staining of a transverse section of gut epithelium from age-matched WT and *olSedl^-/-^* medaka. Note the multiple layers and lumen reduction in *olSedl^-/-^*. Magnification: 63x. Data information: (D) WT n = 50, *olSedl^-/-^* n = 31. *p = 0.02, two-tailed unpaired Mann-Whitney test. Scale bars in (A) and (B): 50 μm; (C): 2 μm; (E): 25 μm.

In addition, we observed an increase of apoptotic cells in retinal tissue of *olSedl^-/-^* at stage 34 (Fig EV4C, D, G), when *olSedl* expression levels peak (Fig 1A) and the highest rate of photoreceptor differentiation in the retina of medaka fish occurs (Kitambi & Malicki, 2008). Notably, at the same stage, *olSedl^-/-^* showed a reduction in proliferating cells, as determined by immunostaining for phosphorylated histone H3 (Fig EV4E, F, H).

Altogether, these data indicate that *olSedl* is required for photoreceptor differentiation and maintenance.

### Sedlin is necessary for intestinal epithelium polarization and development

As noted above, genes belonging to the functional class intestinal absorption were significantly downregulated both in MO_Sedl_ and *olSedl^-/-^.* These genes belonged mainly to the cell component class brush border and included solute and fat transporters (e.g. SLC26A6, SLC3A1, CD36, NPC1L1) and regulatory proteins (PDZK1) present on the apical membrane of intestinal cells, together with structural components of the microvilli (EZR, ESPN). These findings prompted us to analyse the intestine of *olSedl^-/-^* fish. To this end, semi-thin (1 μm) transverse sections at a comparable cranio-caudal axis level were obtained from both WT and *olSedl^-/-^* medaka embryos and stained with toluidine blue. Morphological analysis of posterior gut sections showed a disorganization of epithelial tissue with alterations of cell polarization and a reduction of epithelial cell layers in *olSedl^-/-^* relative to age-matched WT embryos (Fig 5E) in which the gut epithelium is a bilayer sheet and has a radially organized structure with basal-side nuclear localization. Furthermore, the elongated shape of the WT gut epithelium cells in the *olSedl^-/-^* embryos is reduced.

Moreover, the structural alteration of the epithelial tissue in *olSedl^-/-^* was associated with a significant reduction of the gut lumen (Fig 5E). These data suggest a central role for Sedlin in key phases of gut morphogenesis.

## Discussion

Eight distinct genetic disorders are caused by mutations of eight different components of the TRAPP complex: four (TRAPPC2, TRAPPC2L, TRAPPC4, TRAPPC6B) belonging to the core TRAPP complex (common to TRAPPII and TRAPPIII), two subunits (TRAPPC9, TRAPPC10) specific for the TRAPPII complex, and two (TRAPPC11 and TRAPPC12) specific for TRAPPIII (Sacher *et al,* 2019). The clinical manifestations of TRAPPC2 mutations that cause SEDT are distinct from those of the remaining seven. SEDT has cartilage-restricted manifestations and the age-of-onset of clinical signs is in puberty while the other seven TRAPPopathies have neurological and muscular manifestations and a much earlier onset (Sacher *et al,* 2019). The cartilage defects are consistent with the role of Sedlin in procollagen transport that we have shown previously in cell systems (Venditti *et al,* 2012) and confirmed here *in vivo*, and with the extent of collagen secretion during chondrogenesis largely exceeding that of other tissues, while the cartilage-restricted phenotype and later onset could be related to differential temporal/spatial expression of the pseudogene TRAPPC2B.

We have reported here the first animal model for SEDT in medaka fish, which has only one Sedlin-encoding gene. The skeletal phenotype (platyspondyly/reduced length) is highly reminiscent of that observed in SEDT patients, confirming that Sedlin plays a major role in skeletal development. The altered deposition of collagen and the dilation of the ER observed in patient tissues were reproduced in Sedlin KO fish. The *olSedl^-/-^* fish not only represent a reliable model for SEDT but have also revealed multiple aspects that help in understanding the role of Sedlin in development, and hence in the pathogenetic basis of the disease. Altered deposition of Col2A at the ECM in the *olSedl^-/-^* fish leads to ECM disorganization and affects expression not only of the collagen II gene but also other collagen genes such as collagens XI and X, the latter a marker for chondrocyte maturation. Defective ECM deposition affects chondrocyte differentiation markers and the expression of ECM components (Wai *et al,* 1998; Barbieri *et al,* 2003; Komori *et al,* 2022) and while cell-matrix interactions impact on chondrocyte differentiation and cartilage development through cell receptors and their signal transduction pathways (Prein & Beier, 2019), how they impact on the expression of genes encoding ECM proteins is not well understood.

In addition to confirming that Sedlin plays a major role in skeletal development, we also observed additional phenotypes in *olSedl^-/-^* that affect the eye and the gut. These may emerge in medaka because there is only one Sedlin-encoding gene and are not observed in humans due to the presence of the expressed pseudogene TRAPPC2B (Ghosh *et al,* 2001; Vinckenbosch *et al,* 2006). These extraskeletal manifestations could be due to altered ECM deposition in the eye or in the gut, which is an important determinant for their correct development. The ECM plays a key role in retinal cell proliferation, homeostasis, and survival (Pouw *et al,* 2021). In addition, a remodeling of various ECM molecules has been associated with retinal neurodegeneration (Reinhard *et al,* 2017; Wiemann *et al,* 2020). Among the ECM components, collagen II plays a pivotal role in eye development as seen by the ocular manifestations that are associated with mutations of PCII, such as Stickler syndrome type I (membranous vitreous type, OMIM:#108300), Stickler syndrome type I (predominantly Ocular, OMIM:#609508), and vitreoretinopathy with phalangeal epiphyseal dysplasia (OMIM: #619248).

A non-mutually exclusive hypothesis, in line with eye phenotypes reported in COPII mutant zebrafish (Schmidt *et al,* 2013), is that Sedlin may be required for rhodopsin trafficking, which occurs at an extremely high rate in photoreceptors. This possibility is supported by the mis-localization of rhodopsin that we observed in rod cells (Fig 5A and Fig EV4A), where rhodopsin can be seen throughout the cell bodies rather than in the outer segments. In addition, the alteration of collagen gene expression in the *olSedl^-/-^* larvae was accompanied by defects in cell death and proliferation, indicating that Sedlin is dispensable for early steps of eye development but has an unanticipated role in photoreceptor differentiation and maintenance.

Finally, another unexpected role for Sedlin was found to be in gut morphogenesis. Although the basis of this defect will require further analysis, it has been reported that mutation of Sec13 in zebrafish, a subunit of the outer coat of the COPII complex, affects organogenesis of the digestive system, including disruption of the ER in chondrocytes (Niu *et al,* 2012). Furthermore, Sar1B loss of function results in chylomicron retention disease, a rare recessive condition with defects in fat malabsorption in hepatocytes and enterocytes (Levy *et al*, 2019). Together, this evidence supports a model in which finely-tuned regulation of the COPII machinery is critical for gut function, corroborating our data.

In summary, we generated and characterized the first vertebrate model for SEDT, which serves as a valuable tool for gaining deeper mechanistic insights into developmental disorders arising from abnormal Col2 transport and extracellular matrix (ECM) deposition due to membrane trafficking defects.

## Materials and Methods

### Medaka fish stocks

Medaka fish (*Oryzia latipes*) from the Cab inbred strain were used throughout the study and maintained following standard conditions (12h/12h dark/light at 27°C). Embryos were staged according to the method proposed by Iwamatsu (2004). All studies on fish were conducted in strict accordance with the institutional guidelines for animal research and approved by the Italian Ministry of Health, Department of Public Health, Animal Health, Nutrition and Food Safety in accordance with the law on animal experimentation (D.Lgs.26/2014). Furthermore, all fish treatments were reviewed and approved in advance by the Ethics Committee at the TIGEM Institute (Pozzuoli (NA), Italy).

### Whole-mount *in situ* hybridization

Whole-mount RNA in situ hybridization was performed and photographed as previously described (Conte & Bovolenta, 2007). A digoxigenin-labelled anti-sense riboprobe for *olSedlin* was used. Probes were generated from cDNA amplified from st40 larvae using specific primers (Appendix Table 1) and cloned into the Topo-TA-vector (Invitrogen). Larvae were sectioned using a Vibratome (Leica) at 25 µm and images were acquired using Leica DM6000 microscopy.

### Real-time qRT-PCR

Transcriptional levels of *olSedl*, *olBet3* and *olCol2a1* were analysed by quantitative real-time RT-PCR. RNAs were obtained from whole mount larvae at the stages indicated in the legend to Fig 1A using a RNeasy Mini Kit (Qiagen, UK), according to the manufacturer’s protocol. 1 µg of total mRNA from each sample was retrotranscribed using QuantiTech reverse Transcription Kit (Qiagen). Real-Time PCR was performed using SYBR Green Master Mix (Bio-Rad) using the primers listed in Appendix Table 1. Each reaction was performed in triplicate using 25 ng of cDNA in 20 μl. The results were normalised against an internal control (*olHprt*).

### Genomic analysis and generation of *olSedl^-/-^* medaka

The medaka *olSedlin* genomic sequence was retrieved from public databases (http://www.ensembl.org/Oryzias_latipes; ENSORLG00000025160). Custom-designed transcription activator-like effector nucleases (TALEN) were used to induce targeted mutagenesis in the exon 3 of the *olSedl* gene. Potential TALEN target sites were identified by using the TALEN Targeter program at (https://tale-nt.cac.cornell.edu/node/add/talen). Custom-designed TALEN vectors were assembled by Zgenebio (Zgenebio Biotech Inc, Taiwan). The left (L) and right (R) recognition sequences and the spacer sequence on the *olSedl* gene are reported in Appendix Table 1. TALEN RNA was synthesized by transcription using a commercial kit (mMESSAGE mMACHINE SP6 Transcription Kit, Ambion, Life Technologies) and injected into fertilized eggs at the 1-2 cell stage, at a concentration of 100 ng/μl. After hatching, 28 larvae were sacrificed and lysed for genomic DNA extraction. The target region on the *olSedl* gene was amplified by PCR using the primers indicated in Appendix Table 1, and then sequenced to determine the presence of TALEN-induced mosaicism. G0 generation fish from the TALEN-injected embryos and G1 heterozygous fish were selected to generate G2 populations. G2 and G3 populations were genotyped through PCR amplification on genomic DNA extracted from the caudal fin (primers are listed in Appendix Table 1). Once amplified, samples were incubated with the restriction enzyme Xmn1 (NEB, USA), which is able to cut only the WT DNA, as the sequence recognized by the enzyme is lost after TALEN-induced mutagenesis (Appendix Fig S1).

### Bone and cartilage staining

For vital staining, 0.01 % Alizarin Red S (ARS, 3,4-Dihydroxy-9,10-dioxo-2-anthracenesulfonic acid sodium salt, Sigma-Aldrich, A5533) was prepared in embryo medium (0.137 M NaCl, 5.4 mM KCl, 0.25 mM Na_2_HPO_4_, 0.44 mM KH_2_PO_4_, 1.3 mM CaCl_2_, 1.0 mM MgSO_4_, 4.2 mM NaHCO_3_). The specimens were kept in the staining solution for 15 min, then washed at least three times in embryo medium. After no longer than 30 min, specimens were anaesthetized and imaged under green (510–550 nm) fluorescent light using a Zeiss LSM 700 microscope.

For cartilage staining, the fixed medaka larvae were incubated in a freshly prepared cartilage staining solution containing 0.02 % alcian blue (Sigma-Aldrich), 0-200 mM MgCl_2_, and 70 % ethanol. Samples were thoroughly washed in 70% ethanol. After the background colouration was removed, the stained medaka larvae were moved through a graded series of glycerol (50 % to 90 %) and then kept in 100 % glycerol. Images were acquired using a Leica DM6000 microscope.

For staining with alcian blue and alizarin red, the alcian blue stained larvae were washed briefly with a solution containing 20 % ethylene glycol and 1 % KOH, then incubated in a freshly prepared bone staining solution containing 0.05 % alizarin red (Sigma-Aldrich), 20 % ethylene glycol, and 1 % KOH at RT for 30 min with gentle agitation. Samples were thoroughly washed in a prewarmed clearing solution containing 20 % polyoxyethylene (20), 0.1 % Tween 20 and 1 % KOH at 42°C for 3 h or more with agitation.

### MO and mRNA injections in medaka embryos

Morpholinos (Gene Tools, LLC Philomath, OR, USA) were designed against the ATG translational start-site on the *olSedl* gene. Sequences are reported in Appendix Table 1.

Fifty picolitres of Mo-Sedlin solution (approximately 1/10 of the cell volume) were injected at a 0.09 mM concentration into one blastomere at the one/two-cell stage as described previously (Conte *et al,* 2010). A morpholino designed against the *Oryzias latipes* p53 gene (*olp53)* gene was used to control the specificity and inhibitory efficiencies of the morpholinos and the absence of off-targeting effect due to activation of p53, as previously described (Conte *et al,* 2010).

### Immunostaining analysis in medaka fish embryos

Fish at stage 40 were subjected to anesthesia and then fixed by incubation in 100 % methanol for 2 h at room temperature (RT). Samples were rinsed three times with PTW 1× (1× PBS, 0.1% Tween, pH 7.3) and then incubated overnight in 15 % sucrose/PTW 1× at 4°C, and then again incubated overnight in 30 % sucrose/PTW 1× at 4°C. Larval cryosections were processed for immunostaining according to the following procedure. They were rehydrated in 1× PBS for 30 min, washed in PBS/0.1 % Triton X-100 and treated with antigen retrieval solution [proteinase K 20 mg/ml (Sigma-Aldrich) dissolved in 10 mM Tris pH 8.0, 1 mM EDTA] for 15 min at 37°C. Cryosections were then permeabilized with 0.5 % Triton X-100 in 1× PBS for 20 min at RT, rinsed in PBS 0.1% Triton X-100 and moved to blocking solution (2% BSA, 2% serum, 2 % DMSO in PBS/0.1 % Triton X-100) for 30 min at RT.

Cryosections were incubated with rabbit anti-collagen type II, dilution 1:400 (Rockland, 600-401-104), mouse anti-rhodopsin dilution 1:400 (Abcam, 4D2), or rabbit anti-Zpr-1 dilution 1:400 (Abcam, EPR7595) antibodies overnight at 4°C, then washed with PBS/0.1 % Triton X-100 and incubated with Alexa-Fluor-secondary antibodies (1:400; ThermoFisher, A21206, A21202, A10037, A10042) for 1 hr at RT. Nuclei were stained with DAPI (1:500). Where indicated, WGA, 1:500 (ThermoFisher, W11261) was used to stain connective tissue.

### Whole mount immunostaining

Embryos from stage 30 onwards were fixed in 4 % paraformaldehyde, 2× PBS and 0.1 % Tween-20. Fixed embryos were detached from the chorion and washed with PTW 1×. Embryos were digested 20 min with 1m μg/ml proteinase K and washed twice with 2mg/ml glycine/PTW 1×. Samples were fixed 20 min in 4 % paraformaldehyde, 2× PBS and 0.1 % Tween-20, washed with PTW 1× and then incubated for 2 h in blocking solution (FBS 1 %/PTW 1×), at room temperature. The proliferation rate was analysed using an anti-phospho-histone H3 monoclonal antibody dilution 1:400 (Cell Signaling, 9701), with a peroxidase-conjugated anti-rabbit antibody (dilution 1:200; Vector Laboratories) followed by diaminobenzidine staining. To prevent pigmentation of the RPE, medaka embryos were incubated with phenylthiourea (PTU, Sigma-Aldrich).

### Image analysis

The images were processed with Fiji (ImageJ, National Institutes of Health (NIH)) software. Brightness and contrast were adjusted with Adobe Photoshop, and figure panels were assembled with Adobe Illustrator. Images in Figs. 2A, 3A, and Fig EV1C were created using BioRender.

### Detection of apoptotic cell death

The extent and distribution of apoptotic cell death was determined by TUNEL, using the In Situ Cell Death Detection Kit, POD (Roche), following the manufacturer’s protocol. TUNEL assay was performed on 20 µm medaka retina cryosections. As negative control to evaluate possible non-specific effects, fixed and permeabilized retina sections were incubated with the reaction mix without TUNEL reaction enzyme. Sections were observed with a Leica DM-6000 microscope and then confocal images were acquired using the LSM710 Zeiss Confocal Microscopy system.

### Electron microscopy analysis

Medaka larvae at stage 40 were fixed using a mixture of 2 % paraformaldehyde and 1 % glutaraldehyde prepared in 0.2 M HEPES buffer (pH 7.4) for 24 h at 4°C. Larvae were then post-fixed. After dehydration the specimens were embedded in epoxy resin and polymerized at 60°C for 72 h. Thin 60 nm sections were cut on a Leica EM UC7 microtome. EM images were acquired from thin sections using a FEI Tecnai-12 electron microscope equipped with a VELETTA CCD digital camera (FEI). To assess the gut epithelium tissue organization, toluidine blue staining (1 %) was performed on semi-thin (1 μm) transverse sections at a comparable cranio-caudal axis level from both WT and *olSedl^-/-^* plastic-embedded medaka larvae and the sections were examined by light microscopy.

### Protein isolation and Western blot analysis

Total protein extracts were obtained from a pool of 5 WT (control), and 5 *olSedl^-/-^* larvae for each experiment. Larvae were sacrificed and immediately processed with a pestle in SDS buffer (10mM Tris-HCl pH 8.0, 0.2 % SDS and protease inhibitor cocktail). Protein concentration was determined using the Bio-Rad protein assay (Bio-Rad). A total of 35 μg protein from each sample was resolved by SDS-PAGE and transferred to a nitrocellulose membrane (GE Healthcare). The following antibodies were used at the indicated dilutions: rabbit anti-collagen type II 1:1,000 (Rockland, 600-401-104), rabbit anti-actin 1:1,000 (Sigma-Aldrich, A2066), rabbit anti-sedlin 1:1,000 (Venditti *et al,* 2012). Proteins were detected with horseradish peroxidase (HRP)-conjugated goat anti-rabbit IgG antibody (1:8,000, Merck Millipore) and visualised with the LiteAblot Plus substrate (Euroclone), according to the manufacturer’s protocol. Images were acquired using the Chemidoc-lt imaging system (Uvitec Cambridge).

### Library preparation and deep sequencing

For RNA-seq analysis, total RNA was extracted from a pool of 3 Mo-Sedl stage 40 embryos and 3 control (WT) stage 40 embryos. Total RNA (500 ng) from each sample was prepared using TruSeq RNA sample prep reagents (Illumina) according to the manufacturer’s instructions. Quality control of library templates was performed using a High Sensitivity DNA Assay kit (Agilent Technologies) on a Bioanalyzer (Agilent Technologies). The Qubit quantification platform (Qubit 2.0 Fluorometer, Life Technologies) was used to normalise samples for the library preparation. Using multiplexing, up to 6 samples were combined into a single lane to yield sufficient coverage. The amplified fragmented cDNA of ∼200 bp in size were sequenced in paired-end mode using the HiSeq 1000 (Illumina) with a read length of 2×100 bp. Each library was loaded at a concentration of 8 pM, which was previously established as optimal. An average yield of ∼4.5 Mb was obtained per sample.

For the QuantSeq3’mRNA-Seq, total RNA was extracted from a pool of 3 *olSedl^-/-^*(*olSedl KO*) stage 40 embryos and 3 control (WT) stage 40 embryos. Total RNA (100 ng) from each sample was prepared using QuantSeq 3’mRNA-Seq Library prep kit (Lexogen) according to the manufacturer’s instructions. RNA was quantified and mixed at 10ng/μl. The amplified fragmented cDNA of 300 bp in size were sequenced in single-end mode using the NovaSeq500 (Illumina) with a read length of 100 bp.

### Computational analysis of deep sequencing data

Sequence reads were trimmed using bbduk software (https://jgi.doe.gov/data-and-tools/bbtools/bb-tools-user-guide/usage-guide/) (bbmap suite 37.31) to remove adapter sequences, poly-A tails and low-quality end bases (regions with average quality below 6) and then aligned on oryLat2 reference sequence using STAR (Dobin *et al,* 2013). The expression levels of genes were determined with htseq-count (Anders *et al,* 2015) using the Gencode v19 gene model (Frankish *et al,* 2019). Differential expression analysis was performed using edgeR (Robinson *et al,* 2010). The threshold for statistical significance chosen was False Discovery Rate (FDR) < 0.05. In detail, in the QuantSeq 3’mRNA-Seq (GEO accession number GSE143538), 1,052 medaka genes were differentially expressed and converted to the corresponding human orthologues (414 genes induced and 638 inhibited). In the RNA-seq analysis (GEO accession number GSE186769), 5,013 medaka genes were differentially expressed and converted to the corresponding human orthologues (1,870 genes induced and 2,135 inhibited). The conversion from Ensemble ID of the medaka fish to the human orthologs was performed using the BioMart browser (Cunningham *et al,* 2022). The comparison of the two datasets identified 695 DEGs in common divided into 254 induced and 399 inhibited in both datasets, the remaining 42 being oppositely regulated.

### mRNA sequencing data analysis

To further determine the biological consequences of these differentially expressed transcripts, gene ontology enrichment analysis (GOEA) and Functional Annotation Clustering analysis were performed on the 254 induced and 399 inhibited in both datasets, separately. The DAVID Bioinformatic Resources (Huang *et al,* 2009) was used, restricting the output to biological process terms (BP), cellular compartment terms (CC), and the ‘Kyoto Encyclopedia of Genes and Genomes’ (KEGG Pathway) analyses (Kanehisa *et al,* 2016) were also performed. The threshold for statistical significance of GOEA was FDR < 0.1 and Enrichment Score ≥ 1.5, while for the KEGG Pathway analyses was FDR < 0.1.

### Statistical analysis

Statistical analyses were performed using GraphPad Prism9 (GraphPad Software) or the R software environment for statistical computing (rstatix R package). To test the normal distribution of the data, Kolmorogov-Smirnov test was performed. When data were normally distributed, the statistical significance of difference in measured variables was determined by two-tailed unpaired t-test with Welch’s correction when appropriate. For not normally distributed variables, non-parametric unpaired two-tailed Mann-Whitney test was used. For the experiment shown in Figs EV4G and EV4H, a Two-way ANOVA with Sidak’s multiple comparisons posthoc test was performed.

## Data availability

The datasets produced in this study are available in the following databases:

QuantSeq-3’-mRNA-Seq data: Gene Expression Omnibus (https://www.ncbi.nlm.nih.gov/geo/query/acc.cgi?acc=GSE143538)

RNA-Seq data: Gene Expression Omnibus (https://www.ncbi.nlm.nih.gov/geo/query/acc.cgi?acc=GSE186769)

## Acknowledgements

We thank Michele Santoro for oligonucleotide design, Margherita Mutarelli for genome sequence retrieval and analysis, and Ela Knapik for support in medaka phenotyping. This work was supported by Telethon (grant TGM16CBDM13), the Italian Association for Cancer Research (grant IG2013_14761), and European Research Council Advanced Investigator grant 670881 (SYSMET), the University of Naples Federico II (grant STAR Plus 2020 linea 1), and the Italian Ministry of University and Research (PRIN, 2020PKLEPN) to MADM.

## Author contributions

Francesca Zappa: Investigation. Daniela Intartaglia: Investigation. Andrea M. Guarino: Investigation. Rossella De Cegli: Investigation. Francesco Salierno: Investigation. Elena Polishchuk: Investigation. Cristina Sorrentino: Investigation. Cathal Wilson: writing—review and editing. Ivan Conte: Supervision; writing—original draft; writing—review. Maria Antonietta De Matteis: Supervision; funding acquisition; investigation; writing—original draft; writing—review and editing.

## Disclosure and competing interest statement

The authors declare that they have no conflict of interest.

## Figure Legends

**Figure EV1.**
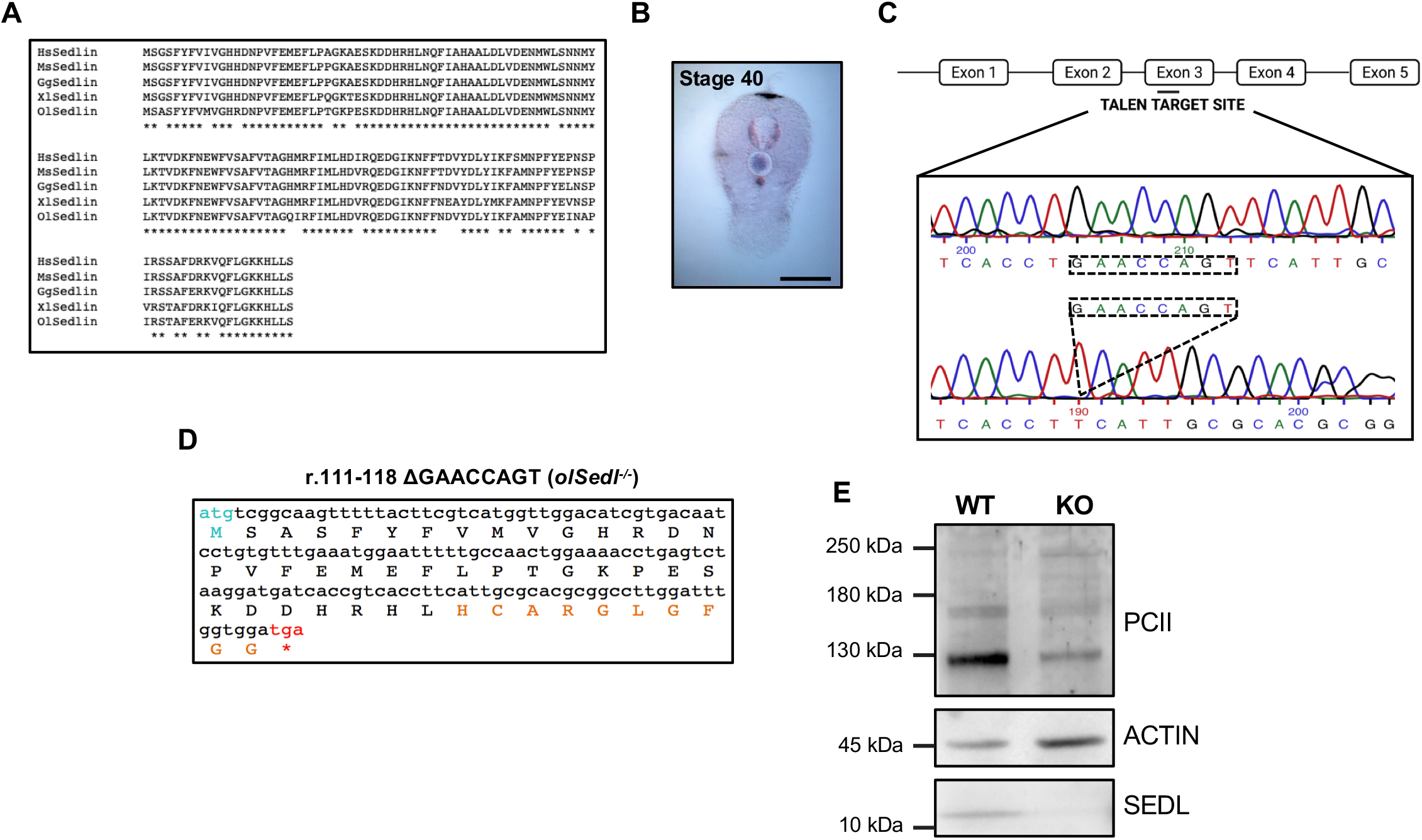
Related to Figure 1: TRAPPC2 gene is conserved in *Orizyas latipes*. A. Multiple alignment of TRAPPC2 (Sedlin) protein sequences. Hs, *Homo sapiens*, Mm, *Mus musculus*, Gg, *Gallus gallus*, Xl, *Xenopus laevis*, Ol, *Orizyas latipes*. B. Representative image of in situ hybridization (ISH) using a DIG-labelled antisense RNA for *olSedl* in vertebrae and muscles of stage 40 larvae. 40x magnification. Scale bar: 1 mm. C. Schematic representation of the *olSedl* gene and the TALEN target site in the third exon. Dashed box in the lower panel indicates the sequence deleted by frameshift mutation. D. Deduced amino acid sequence of the r.111-118 ΔGAACCAGT protein product. The start ATG is shown in light blue, amino acids affected by the frameshift mutation are reported in orange, and the premature stop codon in red. E. Western blot analysis of total lysates from WT and *olSedl*^-/-^ fish probed with anti-Sedlin and anti-PCII antibodies. Anti-actin was used as loading control. Representative image of 2 replicates.

**Figure EV2.**
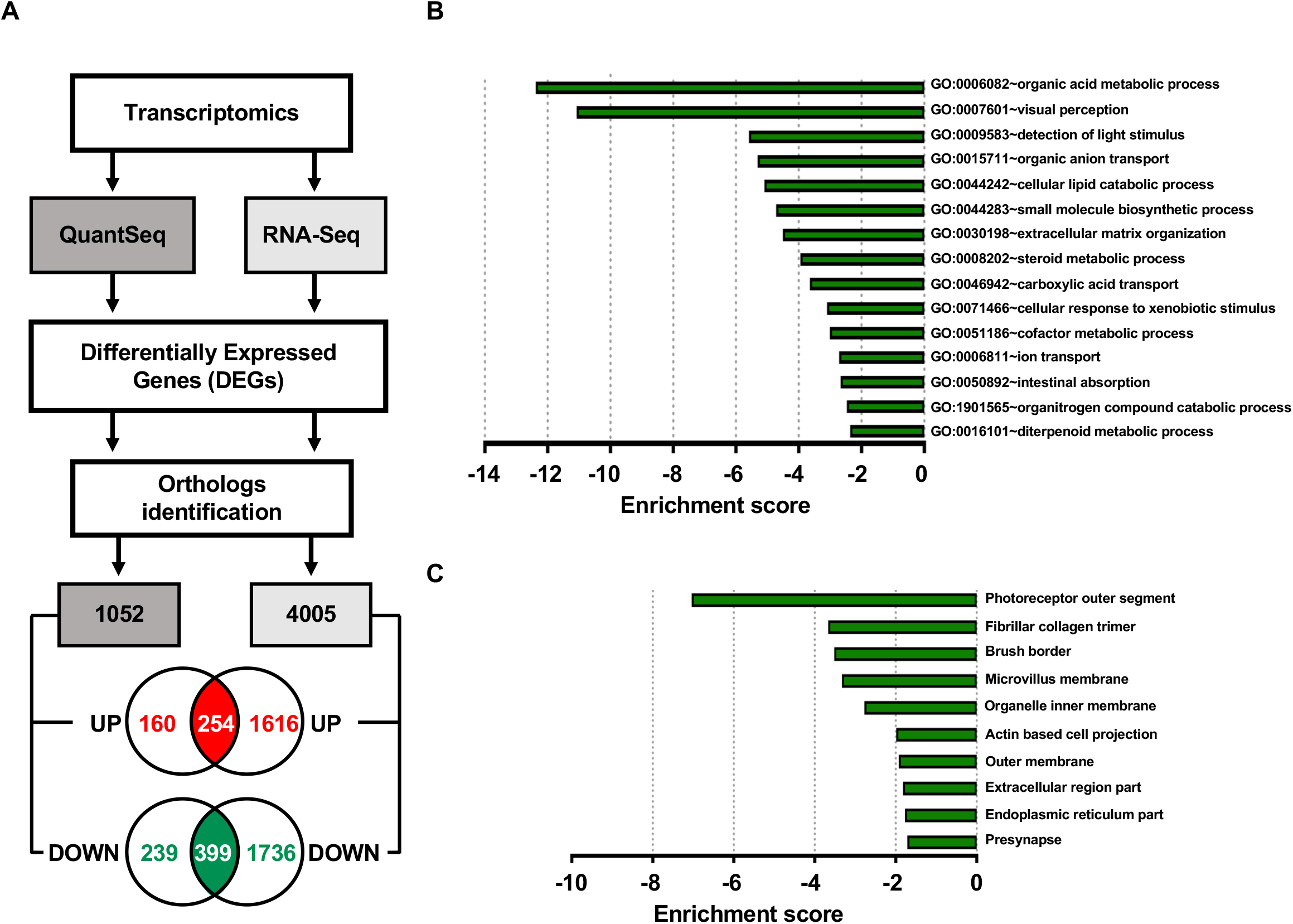
Related to Figure 4: Transcriptome changes induced by *olSedl* depletion. **A.** Schematic representation of QuantSeq-3’-mRNA-Seq and RNA-Seq protocols. For both procedures, differentially expressed genes (DEGs) were converted to the corresponding human orthologs using the BioMart browser (Cunningham *et al,* 2022). The Venn diagrams show the number of up- and downregulated genes for each dataset with 254 upregulated (red) and 399 downregulated (green) common to both datasets. **B.** Bar plots indicating the most significantly enriched Biological Processes terms for the downregulated DEGs common to both datasets. **C.** Bar plots showing the top 10 significantly enriched Cellular Component for the downregulated DEGs common to both datasets.

**Figure EV3.**
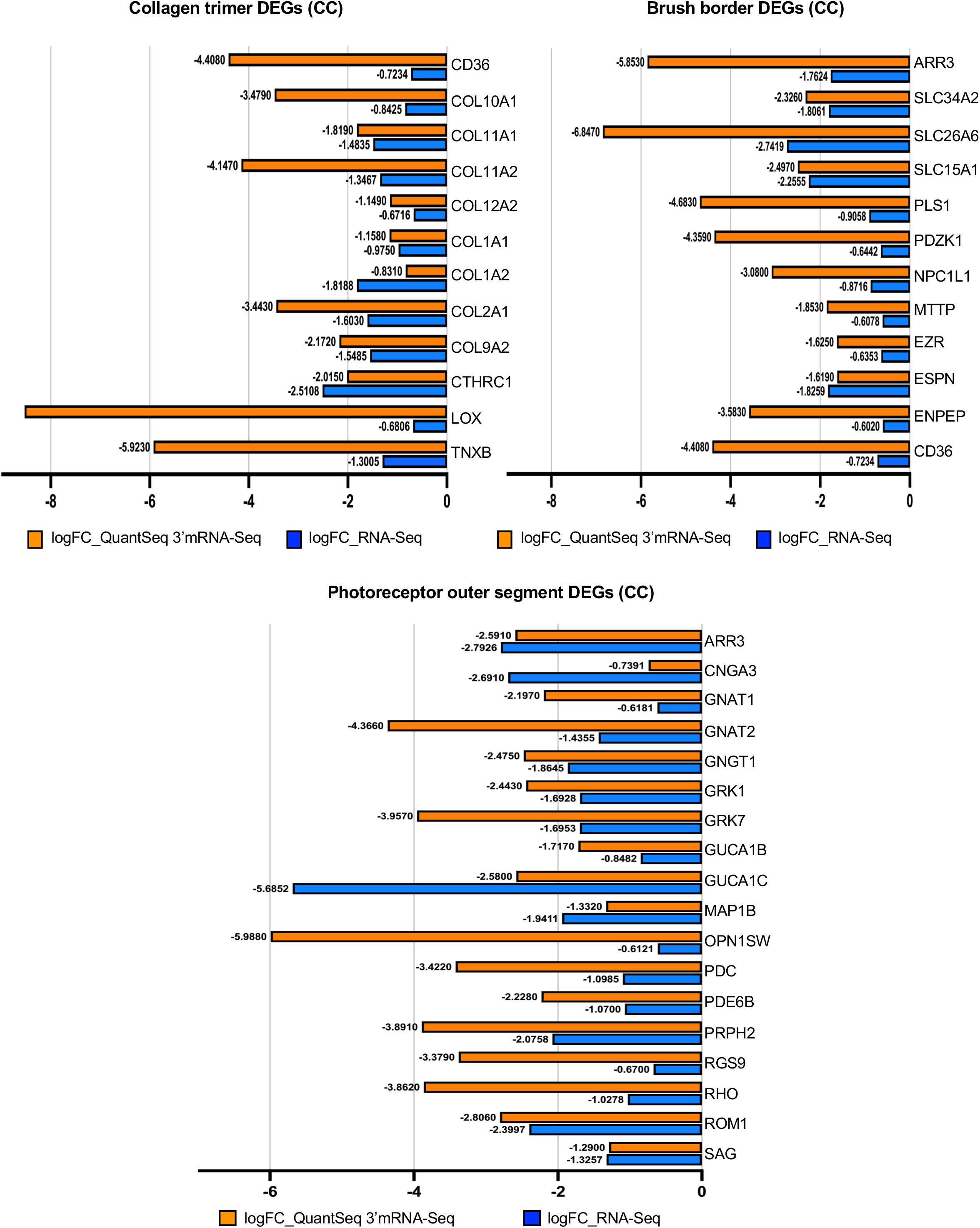
Related to Figure 4: Transcriptome changes induced by Sedlin depletion. Graphical representation of differentially expressed genes (DEGs) in the CC (Cellular Compartments) classes *GO:0005581∼collagen trimer, GO:0001750∼photoreceptor outer segment, and GO:0005903∼brush border* significantly downregulated in both QuantSeq-3’-mRNA-Seq and RNA-Seq datasets.

**Figure EV4.**
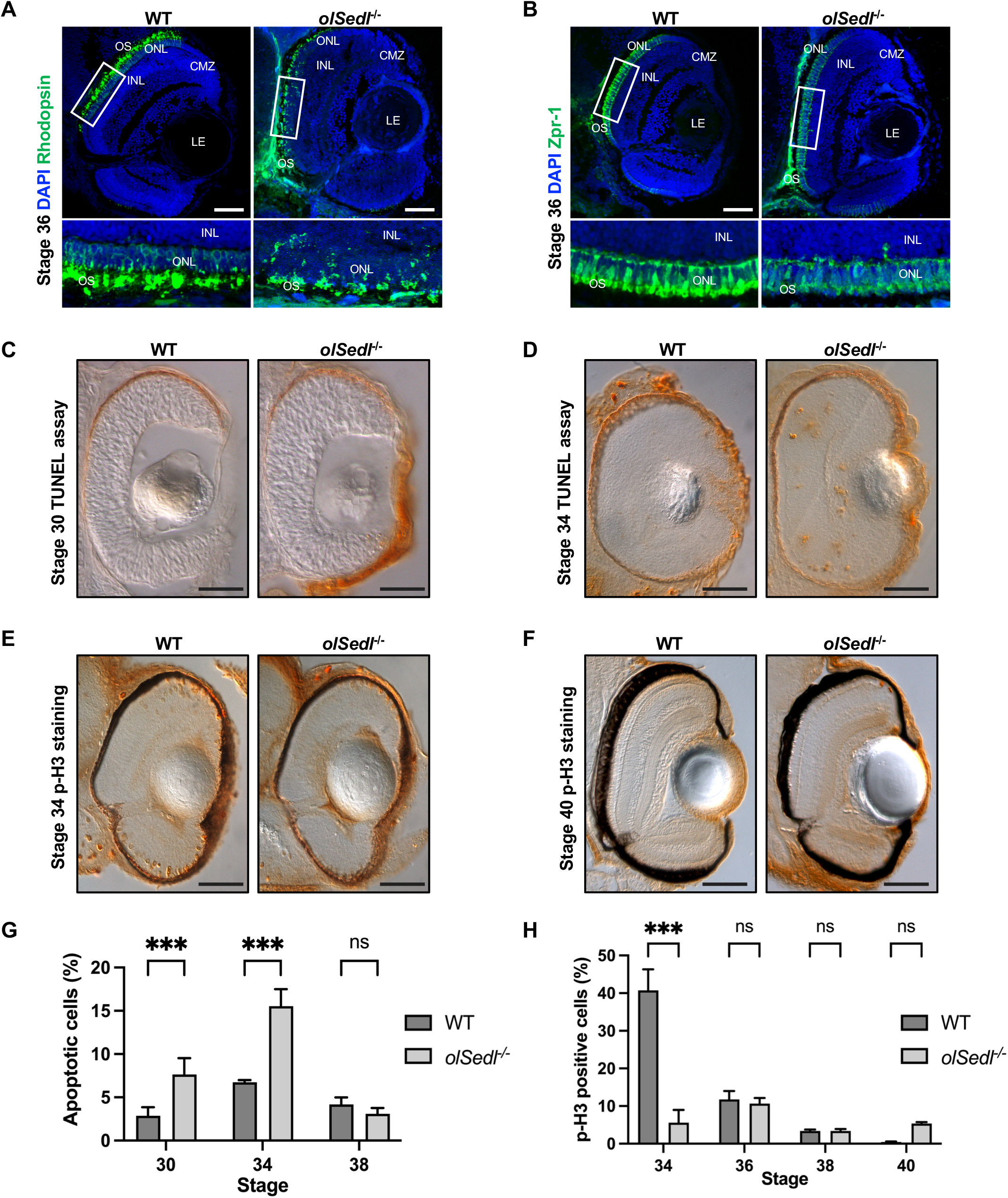
Related to Figure 5: *olSedl^-/-^* fish manifest altered photoreceptor development. A, B. Representative images of Rhodopsin (A, green) and Zpr-1 (B, green) immunostaining of frozen retinal sections of WT and *olSedl^-/-^* stage 36 individuals. Nuclei are counterstained with DAPI (blue). Lower panels are enlarged images of boxed areas [(CMZ) ciliary marginal zone, (OS) outer segment, (IS) inner segment, (ONL) outer nuclear layer, (INL) inner nuclear layer] C, D. Representative images of deoxynucleotidyl transferase dUTP nick end labelling (TUNEL) assay on retinal sections of WT and *olSedl^-/-^* stage 30 (C) and stage 34 (D) individuals. E, F. Representative images of frontal vibratome sections of WT and *olSedl^-/-^* stage 34 (E) and stage 40 (F) individuals immunostained with anti-pH3 (phospho-histone) antibody. G. Quantification of percentage of apoptotic cells at the indicated embryonic stages. H. Quantification of pH3-positive cells at the indicated embryonic stages. Data information: (G) and (H), Mean ± SD, n ≥ 3. ***p < 0.001, two-way ANOVA with Sidak’s post-hoc test. ns, not significant. Scale bars in (A-F): 50 μm.

